# Docclustering: An Implementation of the Novel Ps-Document Clustering Algorithm a Preprint

**DOI:** 10.1101/397133

**Authors:** Jens Dörpinghaus, Sebastian Schaaf, Marc Jacobs

## Abstract

Document clustering is widely used in science for data retrieval and organisation. DocClustering is developed to include and use a novel algorithm called PS-Document Clustering that has been first introduced in 2017. This method combines approaches of graph theory with state of the art NLP-technologies. This new heuristic has been shown to be superior to conventional algorithms and it provides – given a suiting similarity measure – a more accurate clustering on biological and medical data.

Since the application is written for research on biomedical literature, interfaces for PubMed and SCAIView are available. In this brief report the source code as well as a short overview about the new features, novel heuristics and approaches are provided.

The software can be obtained from the authors or directly downloaded from GitHub, see https://github.com/jd-s/DocClustering.

## 1 Introduction

In science, document clustering is widely used for data retrieval and organization as a general technique in text and data mining. It is used to assign documents to clusters, which usually represent topics. In [1] a novel approach using graph theory to document clustering is presented by Dörpinghaus *et al* and discussed together with its application on a real-world data set retrieved from the PubMed database. It is shown that this method is superior to conventional algorithms and provides more accurate clustering on biological and medical data, depending on the chosen similarity measure.

In a nutshell, this method shows that the well-known graph partition of documents to stable sets or cliques can be generalized to pseudostable sets or pseudocliques, which is, for example, described by Schaeffer in [2]. This allows soft clustering as well as hard clustering to be carried out. The presented integer linear programming as well as the greedy approach to this 𝒩 𝒫-complete problem lead to valuable results on random instances and some real world data using different similarity measures. This approach using graph theory comes with two important features: 1) it opens the toolbox of graph theory to the world of clustering and 2) makes it possible to gain a fast but exact access to soft clustering. Basic implementation offers the possibility to cluster data with several similarity measures, given the bounds *∊* and *ι*. In addition, it is possible to use a divide and conquer approach to parallelize the computation and speed up the runtime for large instances.

*DocClustering* as the implemented software package allows application of this new method, that is the principles of soft or hard clustering to documents. *Hard clustering* defines that every document belongs to only one cluster, whereas *soft clustering* allows documents to be classified in one or more clusters. This can be interpreted as probability or as an overlap between topics. Hard clustering produces *stable sets*. In these sets no two nodes are connected. Soft clustering produces *pseudostable sets*. This allows documents to have a position between stable sets. The application naturally suits data obtained from SCAIView or PubMed database, but can be applied to any other data.

Using a graph partition for clustering is widely discussed in literature. Schaeffer points out that “the field of graph clustering has grown quite popular and the number of published proposals for clustering algorithms as well as reported applications is high” [2]. Usually directed or weighted graphs are the subject of research. However, this paper points out that for problem complexity reasons it is suitable to focus on simple graphs. The work reported in [3] explains that a graph partition in cliques or stable sets is most common. Following Jain *et al*, this new approach splits the process into two steps [4]. At first, a similarity measure appropriate to the data domain needs to be defined. Then, the technical clustering process can be performed using an approach of graph theory. Jain *et al* also suggest a final step called “assessment of output” and it is shown that this can also be solved using graph theory and building the graph visualization proposed in [1].

Few authors, like Stanchev in [5], use graph-based approaches. Some authors, e.g. Hirsch *et al*, cover related problems like clustering in the context of search queries [6], whereas Lee *et al* work on the field of hierarchical clusterings [7]. Thus, it can be concluded that focusing only on graph clustering is a novel approach and the generalization of soft document clustering introduced in [1] provides new insights into document clustering – or clustering in general – from the developed graph theoretical toolbox.

DocClustering was implemented using Python 3 and especially the packages `networkx, nltk, sklearn` and `pulp`. DocClustering has two operation modes: A command line mode and an interactive mode that is available via iPython or Jupiter Notebook. Currently, all important features are available in the interactive mode. The software pipeline presented here has three steps: reading data, doing the clustering and evaluating the output (see Figure 1). In interactive mode all parameters can be changed during runtime, whereas in command line mode all parameters have to be set before executing. The amount of PMIDs the tool can handle is limited by main memory only. More than 21,000 documents have been successfully processed in a single run on a computer with a 48GB memory. Here, calculating the similarities is the bottleneck, the clustering process itself only takes a few minutes. In the next section, the clustering approach is described the input and output steps are discussed later in the paper.

**Figure 1:**
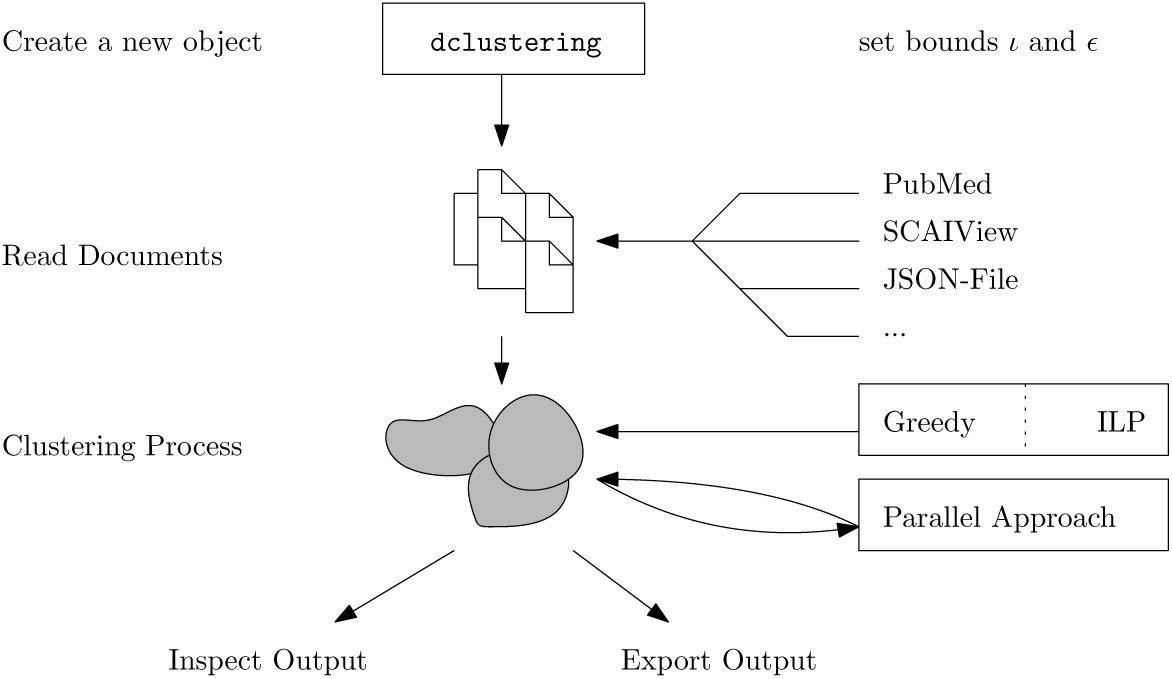
The working process of the application. Reading data, running Greedy or ILP, evaluation of the output.

## 2 Background

In this section, the background of the algorithms used to solve the clustering process is briefly described. For a more detailed overview of the approaches please see [1]. The reading process reads all relevant data and computes the document graph *G* where each node represents a document. As a starting point, the definition of two bounds *ι* and *∊* is required. Since the similarity measure returns values in [0, 1], a bound *∊* need to be defined that separates similar and dissimilar documents. Another bound set is *ι*, with *ι ≥∈*, which is used to describe documents that are not similar enough to be assigned to a cluster but share sufficient similarity to be connected to it. Thus, if *ι* = *∊* is set, there is hard document clustering. Choosing the parameters *ι* and *∊* can be supported by experimental evidence. Further research needs to be done to prove concepts. Usually values between 10% and 40% lead to a reasonable and valuable output.

### 2.1 Integer Linear Program

The Integer Linear Program Approach, introduced in [1] and implemented in DocClustering, can calculate optimal solutions of the multiple pseudostable sets partitioning problem (*minMPS’-a-IP* as given in [1]). It defines constraints to all relevant steps of the underlying graph partition and has exponential runtime. In our experimental results, it only returns output in acceptable time for instances with less than 30 documents. Thus, the purpose of this approach can only usually be to compare the results of heuristics to the optimal solution.

### 2.2 Greedy-Approach

The Greedy-Approach, which is also introduced in [1] and which is made available in DocClustering, can solve the minMPS’-a-IP problem in polynomial time and provides sufficient, but not necessarily global optimal solutions. This is due to the fact that the problem is 𝒩 𝒫 -complete. It starts with an approximation of a graph coloring, leading to a graph partition into stable sets. Starting with the smallest stable set, it attempts to eliminate all possible documents in this set and find an optimal local solution. As a result, small clusters are removed and transformed into documents lying between the clusters. In [1], the following heuristics to start the graph coloring are used: the *greedy independent sets* (GIS) approach [8], the SLF approach (see [8] and [9]) and the Clique Partition on 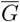^1^ by applying the TSENG clique-partitioning algorithm [10]. All of them are available through the methods `runSLF, runGIS` and `runClique`. The SLF approach is the recommended heuristic for this application, providing valuable output within an acceptable time. Currently, the derivation of a function *p*, which will allow the conversion of this greedy approach into a *p* approximation algorithm, is subject to ongoing research. At this point, it can only be said that the solution is better than the heuristic used for graph coloring and worse than the optimal solution.

## 3 Implementation

The command line tool is run from `clustering.py`. It outputs `.gml`, `.csv` and `.json` files for all heuristics to a destination folder. The logging output is printed to the screen. A default configuration is set up in `etc/config.cfg` and can be modified as required. For an exhaustive list of parameters we refer to the documentation.

For the interactive variant, all clustering data are stored in a dclustering object:

~~~
c l u s t e r = d c l u s t e r i n g ()
~~~

When creating this new object, the optional similarity bounds *∊* and *ι* can be set (e.g. `cluster = dclustering(0.1, 0.4)`; see Section 2 for details) or left to defaults.

DocClustering incorporates readers that can handle different input formats. `read_PmidList` for example allows to read a list of PMIDs, whereas `read_SCAIview` reads the `json-`formatted export from SCAIView. An example is:

~~~
c l u s t e r . r e a d F r o m F i l e (“ / tmp / Alzheimers_ Dement_ pmids . j s o n ”)
~~~

The document graph is automatically built with the selected bounds. It is also possible to specify the similarity measure, since DocClustering includes a large set of build-in measures. For instance, `sim_Mesh` computes the Jacard distance according to the MeSH terms, and `sim_text` computes the TF.IDF measure on the abstract or – if available – the full text. Notably, the choice of similarity has a deep impact on the results. The clustering will either automatically run as soon as you use the methods for feature selection or can be run manually with the functions mentioned in Section 2.2.

The built-in functions can either be used for feature selection and/or output analyses, for example:

~~~
c l u s t e r . g e t C l u s t e r s ()
c l u s t e r . g e t C l u s t e r (8)
~~~

The first method returns a complete list of all clusters, whereas the getCluster() method calculates the content of a distinct cluster as dictionary, containing summed MeSH-terms (topics), PMIDs (ids), the number of documents in the cluster (value) and the list of documents per year (yXXXX). An example of a shortened output is:

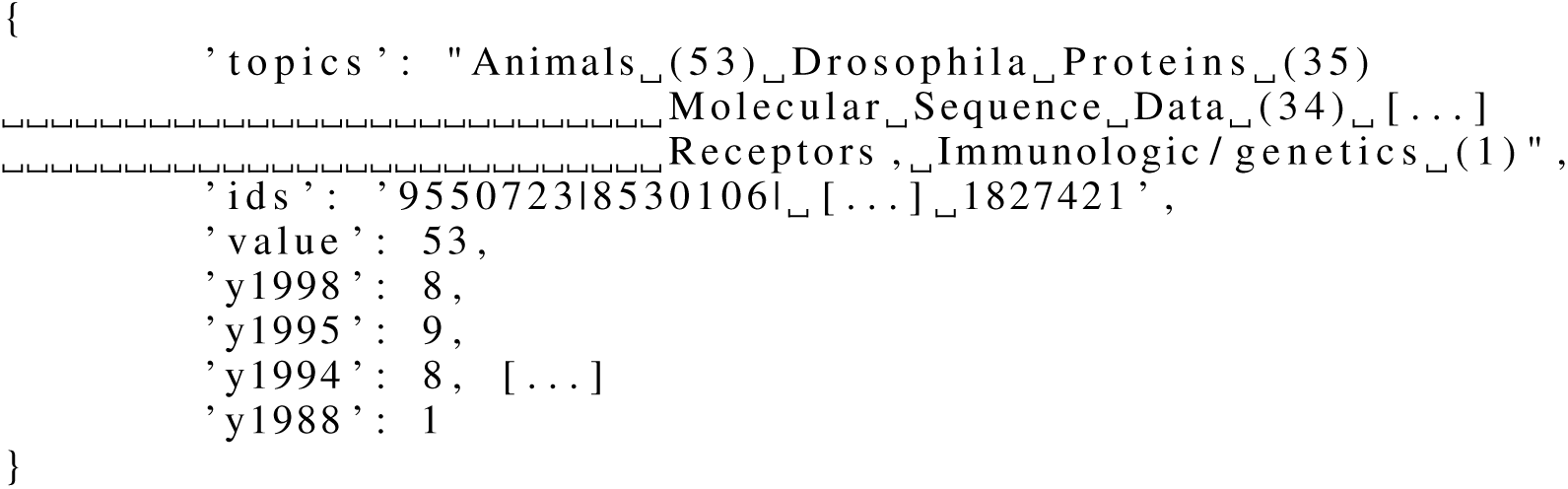

To acquire an overview of clusters and their content, the complete instance graph can be saved as a `.gml-`file for later processing with possibly other tools, for instance Cytoscape, using the following command:

~~~
c l u s t e r . s a v e I n s t a n c e G r a p h (“ / tmp / graph . gml ”)
~~~

## 4. Discussion and Conclusion

This work provides a new Python application “DocClustering”, which is an implementation of novel heuristics and algorithms to perform soft document clustering. An example output of this application can be seen in Figure 2. Further and more detailed results are available in [1].

**Figure 2:**
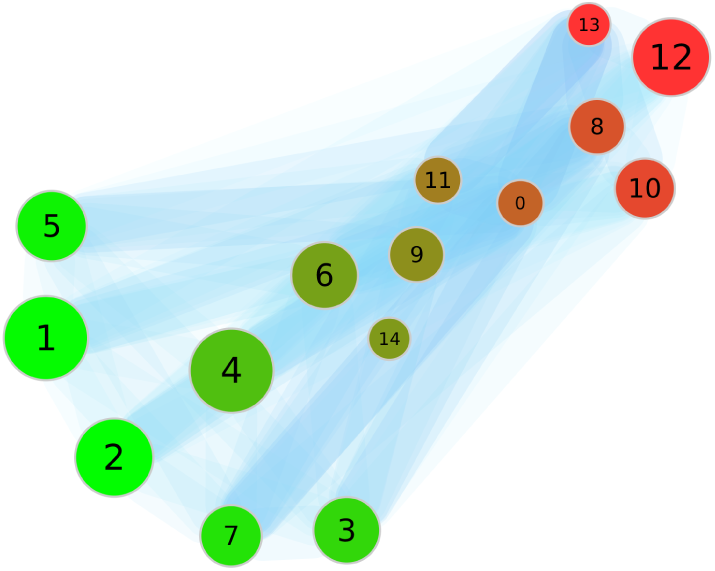
The output of the clustering process of two search queries from PubMed using the SLF approach. Both queries can be separated and are described by the node’s color.

The implemented software can be both beneficial for information retrieval on biomedical documents itself as well as for research on the novel graph theoretical foundation of soft document clustering. While document clustering is employed in numerous text-retrieval applications, this new approach provides a soft clustering with flexible bounds and a solid underlying graph theory heuristic.

DocClustering is platform-independent, open-source, and available for download from GitHub (https://github.com/jd-s/DocClustering). The tool can be executed on any operating system which runs Python Version 3 or higher. The integer linear program also requires a running GLPK installation. The complete documentation of all available functions with some examples can also be found in the Git repository. Computations and visualizations made with DocClustering are available in [1]. Currently, research is being carried out on a better evaluation technique and parallelization of code execution, with the aim of improving overall performance.

## Acknowledgements

Valuable suggestions during the development of DocClustering were provided by Juliane Fluck and Sumit Madan from Fraunhofer SCAI. We would also like to thank the anonymous reviewers for their suggestions that greatly improved the quality of this manuscript.

## Ethics approval and consent to participate

Not applicable.

## Consent for publication

Not applicable.

## Availability of data and materials

DocClustering is Python-based, platform-independent, open source, and can be downloaded from GitHub, see https://github.com/jd-s/DocClustering.

## Authors’ contributions

This new approach goes back to an initial idea of MJ and was developed by JD. The datasets for evaluation were produced by MJ. The design of evaluation process was done by SS and done by all three. TB and JD wrote the manuscript. All authors read and approved the final manuscript.

## Footnotes

1 where 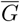 is the complement of graph *G* = (*V, E*), with *V* being the set of nodes and *E* the set of edges

